# The human pathome shows sex specific aging patterns post-development

**DOI:** 10.1101/2023.02.27.530179

**Authors:** Michael Ben Ezra, Jonas Bach Garbrecht, Nasya Rasmussen, Indra Heckenbach, Michael A. Petr, Daniela Bakula, Laust Mortensen, Morten Scheibye-Knudsen

**Affiliations:** Center for Healthy Aging, Department of Cellular and Molecular Medicine, University of Copenhagen; Methods and Analysis, Statistics Denmark; Tracked.bio

## Abstract

Little is known about tissue specific changes that occur with aging in humans. Using the description of 33 million histological samples we extract thousands of age- and mortality-associated features from text narratives that we call The Human Pathome (pathoage.com). Notably, we can broadly determine when pathological aging starts, indicating a sexual dimorphism with females aging earlier but slower and males aging later but faster. Using machine learning, we employ unsupervised topic-modelling to identify terms and themes that predict age and mortality. As a proof of principle, we cross reference these terms in PubMed to identify nintedanib as a potential aging intervention and show that nintedanib reduces markers of cellular senescence, reduces pro-fibrotic gene pathways in senescent cells and extends the lifespan of fruit flies. Our findings pave the way for expanded exploitation of population datasets towards discovery of novel aging interventions.

## Introduction

Aging is a complex, multifactorial process^1,2^ that leads to declining physiology and a susceptibility to disease^3^. Yet, little is known about tissue specific changes that occur with aging in humans. Clinical text constitutes the most abundant data type in electronic health care records which are implemented in most countries^4^. Specifically, pathology records are rich in descriptions of cellular and histological samples of healthy and diseased human tissue and therefore represent a considerable opportunity to systematically characterize tissue specific changes that occur in aging. Nonetheless, electronic health care records are a vastly underused data resource due to their limited availability to researchers^5^. Furthermore, unstructured text data are not directly amenable to computational analysis and clinical text is highly heterogeneous. Importantly, using natural language processing and machine learning, phenotypes can be extracted from clinical text and used to discover correlations and stratify patient cohorts^6,7^.

To get an unbiased description of organismal- and tissue-specific aging, we analyzed the Danish pathology register containing the clinical description of over 33 million samples collected since 1970^8^ from over 4.9 million individuals some born as early as 1876 (Fig. 1a). Using natural language processing we extracted thousands of clinical features from unstructured pathology narrative texts. We combine this with vital statistics to identify age- and mortality-associated features. Using supervised and unsupervised machine-learning we identify population-based patterns of aging and surprisingly discover that pathological aging starts almost immediately after development in the late teens for females. For males, pathological aging starts later (∼40 years) but progresses faster. Conversely, tissue-specific patterns of aging show that some tissues age linearly and others age along developmental and pathological aging trajectories. To further investigate the meaning of clinical features we employed topic-modelling^9^ and reveal specific age- and mortality associated themes. As a proof of principle, we deploy this in lung pathology records and find that the predicative power of topic modelling themes is stronger than individual features. We further cross-reference the age-associated terms from the pathology datasets within all published PubMed abstracts and identify compounds enriched in aging terms. Among them, we identify nintedanib, a tyrosine kinase inhibitor^10^, as a potential pharmacological intervention in aging. Indeed, nintedanib, an antifibrotic agent, reduces markers of cellular senescence, reduces pro-fibrotic gene pathways in senescent cells and extend the lifespan of *drosophila melanogaster*.

**Fig. 1.**
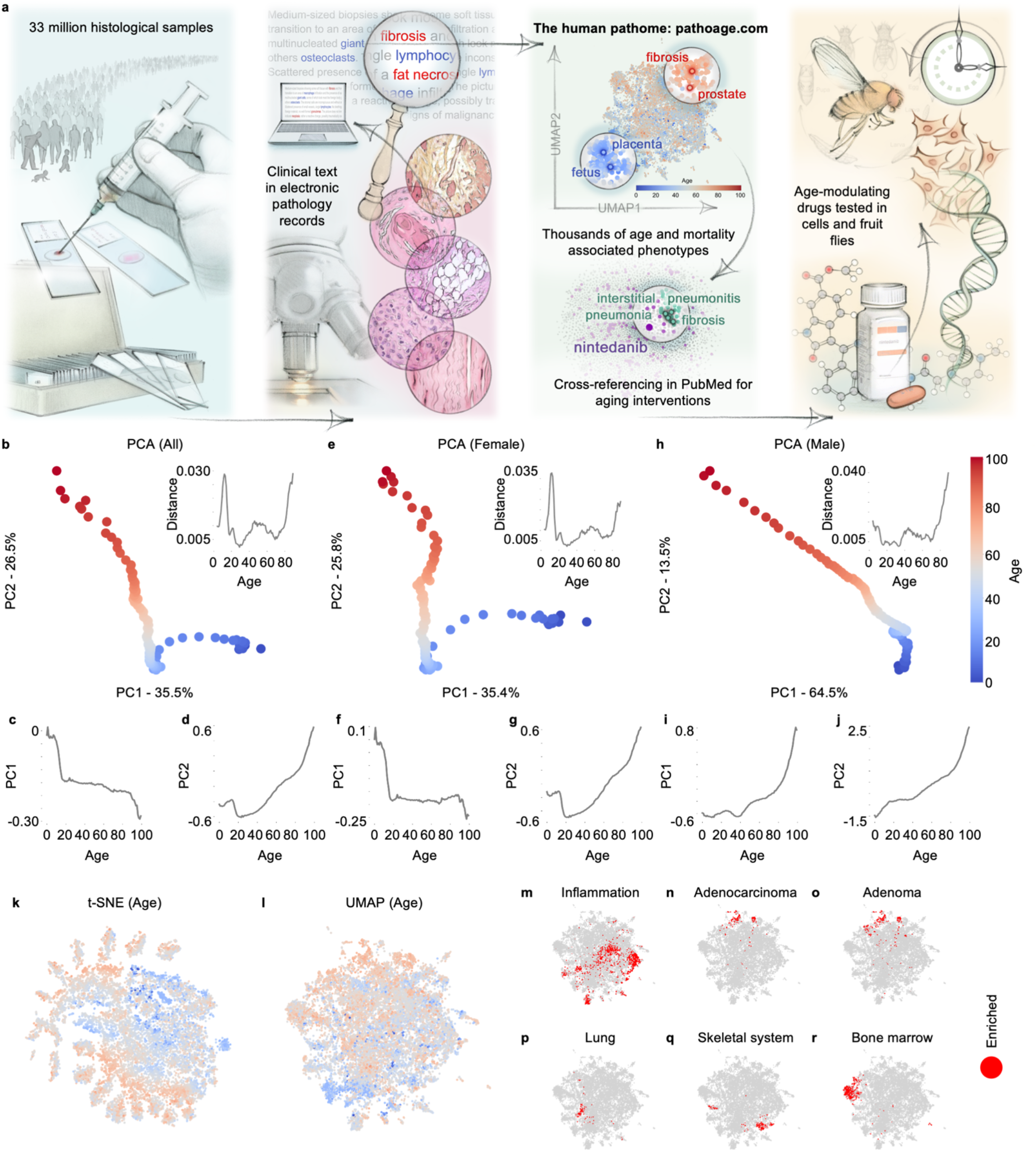
Pathological aging begins post-development for females and at mid-life for males. **a**, The Human Aging Pathome and aging intervention discovery concept and workflow. **b**, PCA of age-aggregated pathology records (n=20,316,270) from in entire pathology register. Normalized Euclidean distance between age adjacent PCA coordinates. **c**,**d**, PC1, PC2 coordinates vs. Age. **e**, PCA of age-aggregated of pathology records (n=14,492,989) from females in the entire pathology register. Normalized Euclidean distance between age adjacent PCA coordinates. **f**,**g**, PC1, PC2 coordinates vs. Age. **h**, PCA of age-aggregated of pathology records (n=5,823,281) from males in the entire pathology register. Normalized Euclidean distance between age adjacent PCA coordinates. **i**,**j**, PC1, PC2 coordinates vs. Age. **k**, t-SNE of clinical features in age-aggregated pathology records in the entire pathology register. **l**, UMAP of clinical features in pathology records in the entire pathology register. **m-o**, Positive morphology specific enrichment: Inflammation, Adenocarcinoma and Adenoma. **p-r**, Positive enrichment of clinical terms in tissue specific pathology records: Lung, Skeletal system and Bone marrow.

## Results

### Pathological aging begins post-development for females and at mid-life for males

To explain the variance in pathology records in the entire pathology register we identified the average term frequency within each age-group (0-100) and performed a principal component analysis. Remarkably, we observed a strong correlation between the main principal components PC1 (35.5%) and PC2 (26.52%) and age, showing that variance in pathology records is strongly explained by age (Fig. 1b). We observed that the ages of development (0-18) vary primarily along PC1 (Fig. 1c) whereas post development ages 19 and over vary primarily along PC2 (Fig. 1d) suggesting that PC1 describes variance in development while PC2 describes true pathological aging-associated changes. We also noted the Euclidean distance of age adjacent PCA components to assess the increase in variance with age. We noted a peak around the end of development, at midlife and late in life. Since we saw increased variance around midlife, we speculated that there could be sex-dependent differences in pathological aging perhaps around menopause. Strikingly, in females age-associated changes appear immediately after development (Fig. 1g) while in males pathological aging appears to start (Fig. 1i) at around forty years of age but does so at an increasing rate. Notably, while aging starts earlier in females the contribution of this factor to the overall variance is much smaller in females than males (PC2 females accounts for 25.8% of variance while PC1 for males account for 64.5%). To visualize the entire landscape of terms, we applied t-distributed stochastic neighbor embedding (t-SNE) (Fig. 1k; Extended Data Fig. 1a,b for sex-specific t-SNEs) to the average term frequencies within each age-group overlaid with the mean incidence age of each feature in the pathology register. Strikingly, the t-SNE visualization shows that terms primarily associated with younger age groups coalesce in the center while terms associated with older-age groups project outwards in all directions reflecting an apparent age-dependent progression from order to disorder.

### Patterns of tissue specific vocabulary identified in pathology records

To visualize the co-occurrence of terms in the entire pathology register we applied Uniform Manifold Approximation and Projection (UMAP). The UMAP shows a more unidirectional age-effect than observed in the t-SNE (Fig. 1l; Extended Data Fig. 1c,d for sex-specific UMAPs). In both t-SNE and UMAP visualizations we observed a tendency for terms with similar mean incidence age to co-occur. Pathology records in the registry are classified according to morphology and tissue (Fig. 1m-r; Extended Data Fig. 1e for additional tissues). We noted that records annotated with the morphology code ‘inflammation’ are enriched with terms from broad clusters of the feature landscape (Fig. 1m). Further, records associated with specific tissues such as lung, skeletal system and bone marrow are enriched in clinical terms from narrower regions of the feature landscape (Fig. 1p-r). Notably, terms enriched in lung tissue coincide with inflammation supporting the notion that inflammation affects lung more strongly compared with some other tissues in the body^11^. In addition, terms enriched in most tissues are largely non-overlapping, suggesting that terms used to describe specific tissues tend to be more distinct. On the other hand, related tissues such as skeletal system (Fig. 1q) and bone marrow (Fig. 1r) are enriched in terms from neighboring regions of the landscape. Altogether, these findings suggest that tissue specific patterns of aging can be identified in the dataset.

### Tissues age along specific trajectories

To identify tissue specific aging patterns, we repeated the above analyses for every tissue in the body. We noted a similarly strong correlation between the two main principal components PC1 and PC2 and age in multiple tissues (Fig. 2a). To understand whether the mean incidence age of terms (Fig. 2b) and the mortality (defined as time to death from examination) associated with terms are correlated, we were able to connect the mortality data of over 1.3 million individuals with their own pathology records (Fig. 2c). Incidentally, hazard of death is associated with the word count length of clinical text narratives and with birth cohort (Extended Data Fig. 1f,g) (Extended Data Fig. 2a-c for additional tissues). Indeed, age and mortality appear broadly correlated in all tissues. Interestingly, in several tissues (kidney, bladder, nervous system) we observe that age-related changes appear to be biphasic (Fig. 2d) perhaps suggesting a phase associated with development and one associated with pathological aging. For other tissues, aging appears more linear (lung, liver, gallbladder, skeletal system, nervous system.) When investigating a single tissue such as lung, clinical features enriched in categories such as inflammation and benign tumors (Fig. 2e,f) appear to be mostly non-overlapping while adenoma and adenocarcinoma morphologies (Fig. 2g,h) appear to coincide. In sum, tissue specific trajectories define different patterns of aging indicating that different tissues age in different ways. To allow exploration of these phenomena, we have created a browsable database of the human pathome (www.pathoage.com).

**Fig. 2.**
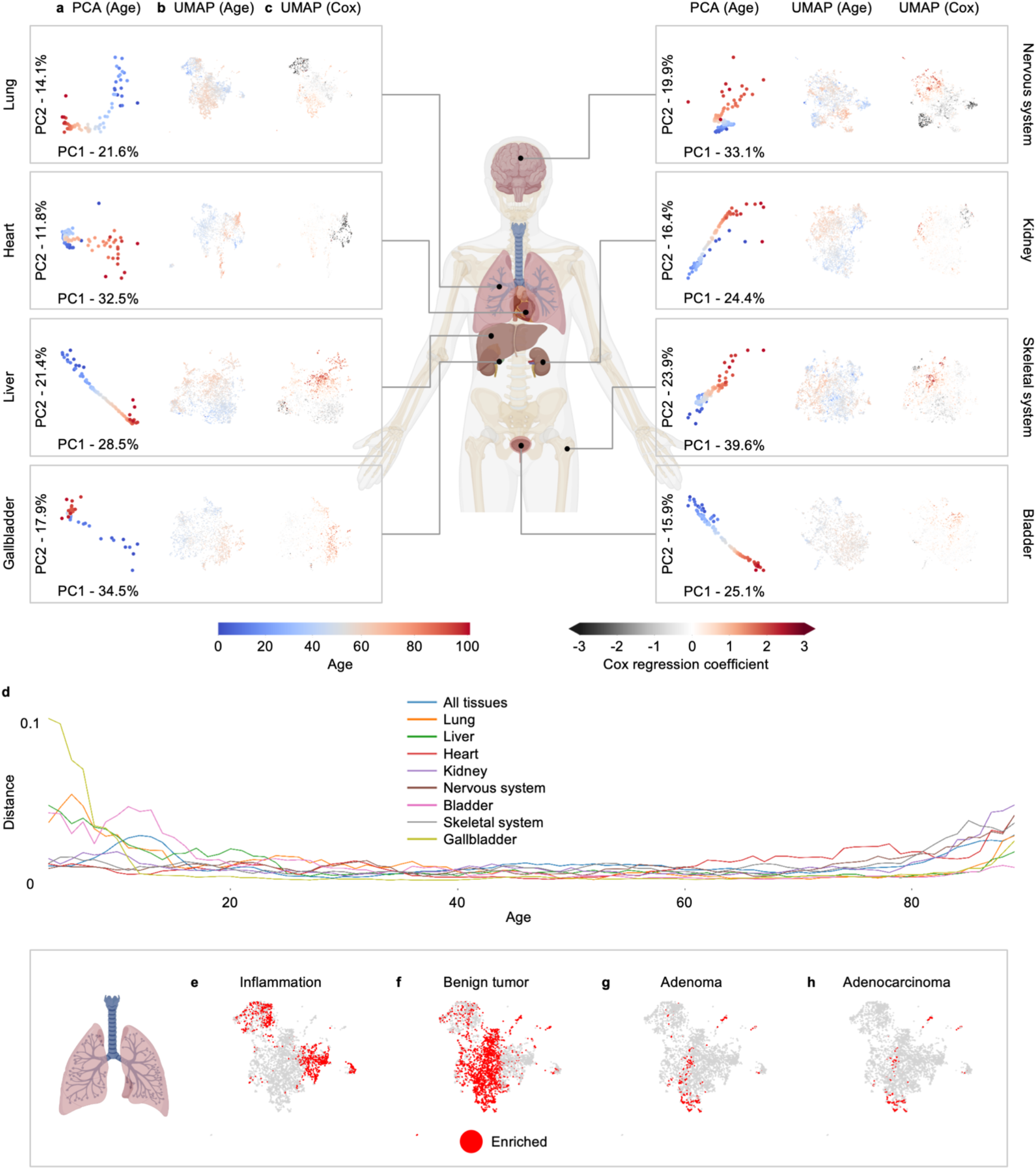
Tissues age along specific trajectories. **a**, PCA of age-aggregated tissues specific pathology records. **b**, UMAP of clinical features of tissue specific pathology records (mean incidence age). **c**, UMAP of clinical features of tissue specific pathology records (Cox regression coefficient).Tissues shown are: Lung (n=177,795), Liver (n=156,057), Heart (n=27,055), Kidney (n=85,244), Nervous system (n=183,729), Bladder (n=250,532), Skeletal system (n=242,282) and Gallbladder (n=182,261). **d**, Normalized Euclidean distance between age-adjacent PCA coordinates of all tissues in. **e-h**, Positive enrichment of clinical terms in morphology specific lung records: **e**, Inflammation. **f**, Benign tumor. **g**, Adenoma. **h**, Adenocarcinoma.

### Clinical features in pathology text predict age

To better understand tissue specific aging, we fit a supervised deep neural network multilayer perceptron regression model to predict age from clinical text features in lung pathology records (Mean absolute error MAE=7.61, Fig. 3a). We performed a feature importance analysis (Fig. 3b) and identified the terms ‘carcinoma’, ‘anthracnose’, ‘planocellular’ and ‘sarcoidosis’ as most predictive of lung aging. However, given the relatively poor predictive power of the model (R^2^=0.29) and the small contribution of each term (ΔR^2^<0.021) we decided to investigate whether a collection of associated terms would yield stronger predictive power.

**Fig. 3.**
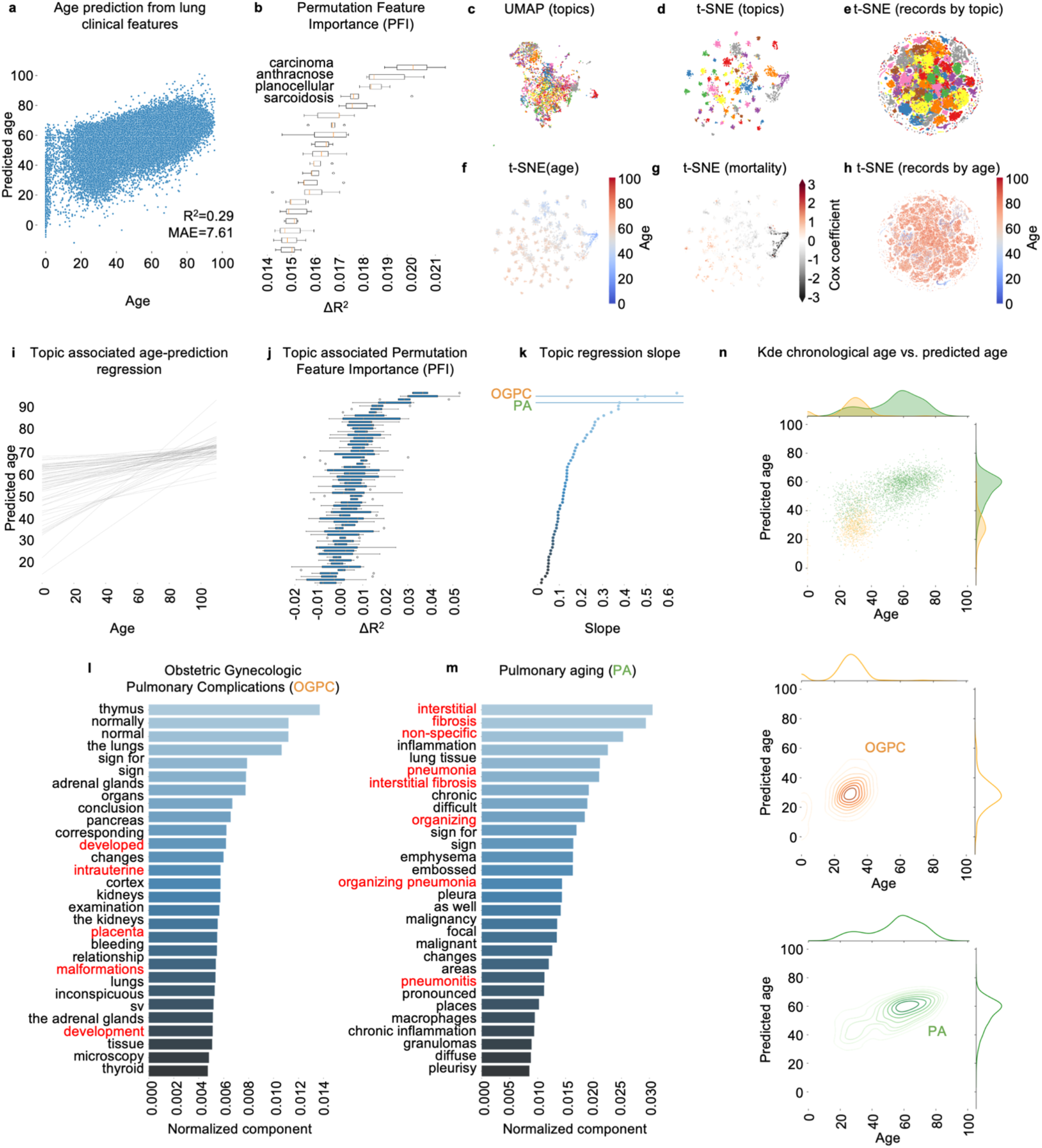
Semantic structures in clinical text describe lung pathologies and predict age. **a**, DNN age-prediction from clinical features in lung pathology records. **b**, Permutation feature importance (PFI). **c**, UMAP of clinical features in lung records (LDA topic). **d**, t-SNE of clinical feature LDA distributions in topics (LDA topics). **e**, t-SNE of LDA record distributions in topics (LDA topics). **f**, t-SNE of clinical feature LDA distributions in topics (mean incidence age). **g**, t-SNE of clinical feature LDA distributions in topics (mortality: Cox regression coefficient). **h**, t-SNE of LDA record distributions in topics (age). **i**, Topic associated age-prediction linear regression. **j**, Topic associated permutation feature importance (PFI). **k**, Topic associated age-prediction regression slope. **l**, Terms describing Obstetric Gynecologic Pulmonary Complications (OGPC). **m**, Terms describing pulmonary aging (PA). **n**, Bivariate kernel density estimation (kde) and histograms of chronological age vs. predicted age, records closely associated with PF and OGPC.

To understand how clinical features semantically relate to one another we applied a latent Dirichlet allocation (LDA) topic model^9^ to tissue-specific pathology records enabling us to identify clusters of co-occurring terms. We applied the topic model to 177,795 lung pathology records and employed a model perplexity minimization strategy to determine that the clinical feature space is optimally decomposed into sixty topics (Extended Data Fig. 3a). A t-SNE visualization of clinical feature distributions in topics demonstrates the topic model’s ability to segregate associated features into clusters (Fig. 3c,d). In turn, this also enables us to stratify individual pathology records by topic (Fig. 3e). Importantly, the topic model appears to identify collections of features with closely associated age and mortality (Fig. 3f,g; Extended Data Fig. 3b,c) suggesting that these semantic structures could describe clinically relevant themes. Importantly, stratified patient records also appear to have closely associated age at examination (Fig. 3h) further strengthening the notion that the topics we identified may characterize cohorts of similar individuals.

### Predicative power of topics is stronger than individual features

To assess the predicative power of collections of associated terms, we performed linear regression on the age and predicted age of records closely associated with each topic (Fig. 3i; Extended Data Fig. 3d). Furthermore, we performed feature importance on the collected terms that make up each topic (Fig. 3j; Extended Data Fig. 3e). Notably, the maximum importance of collections of terms (ΔR^2^<0.039) to age prediction is approximately 2-fold greater than that of the maximum importance of an individual feature (ΔR^2^<0.021).

We then identified topics with changes in the age-prediction regression slope as topics where aging elicits alterations in the age-effect (Fig. 3k; Extended Data Fig. 3f). Among the topics (Supplementary Table 2 for full list) we identified clinical themes consisting of terms broadly describing cases of lung pathologies. One topic appeared to be associated with Human Immunodeficiency Virus (HIV) (terms such as ‘fungi’, ‘pneumocystis’, ‘carinii’, ‘pneumocystis carinii’, ‘alveolar’, ‘inflammation’ and ‘fibrosis’) (Extended Data Fig. 3g) and another with obstetric gynecologic pulmonary complications (OGPC) (terms such as ‘development’, ‘intrauterine’, ‘placenta’ and ‘malformations’) (Fig. 3l). Interestingly, a topic appeared to describe pulmonary aging (PA) with terms such as ‘interstitial’, ‘fibrosis’, ‘non-specific’, ‘pneumonia’, ‘interstitial fibrosis’, ‘pneumonitis’ and ‘fibroelastosis’ (Fig. 3m). Further illustrating the relationship between the age and predicted age of records are kernel density estimation plots corresponding to each of the topic-specific regressions (Fig. 3n; Extended Data Fig. 3h). Notably, the pulmonary aging topic regression shows strong age dependency. In sum, our approach effectively leads us to identify an associated collection of aging modifiers.

### Cross-validation with PubMed identifies age-modifying drugs

Since we had identified terms in the pathology register that are age-associated we could identify any other terms (terms, genes, drugs etc.) in other text-based databases (e.g. PubMed, OMIM.org etc.) that co-occur with these age-associated pathology terms. We decided to investigate molecules that are co-mentioned in PubMed abstracts with clinical terms from our identified lung-aging topic (Fig. 4a). Approximately 35 million molecules in the PubChem library were mined in over 31.8 million PubMed abstracts and assigned a proximity score (Fig. 4b). Among the terms scoring highest we identified nintedanib, a tyrosine kinase inhibitor, as a potential pharmacological intervention in aging. Nintedanib is an anti-fibrotic drug used in the treatment of idiopathic pulmonary fibrosis^12^. Alongside nintedanib we tested axitinib, another tyrosine kinase inhibitor with potential anti-fibrotic effects^13^.

**Fig. 4.**
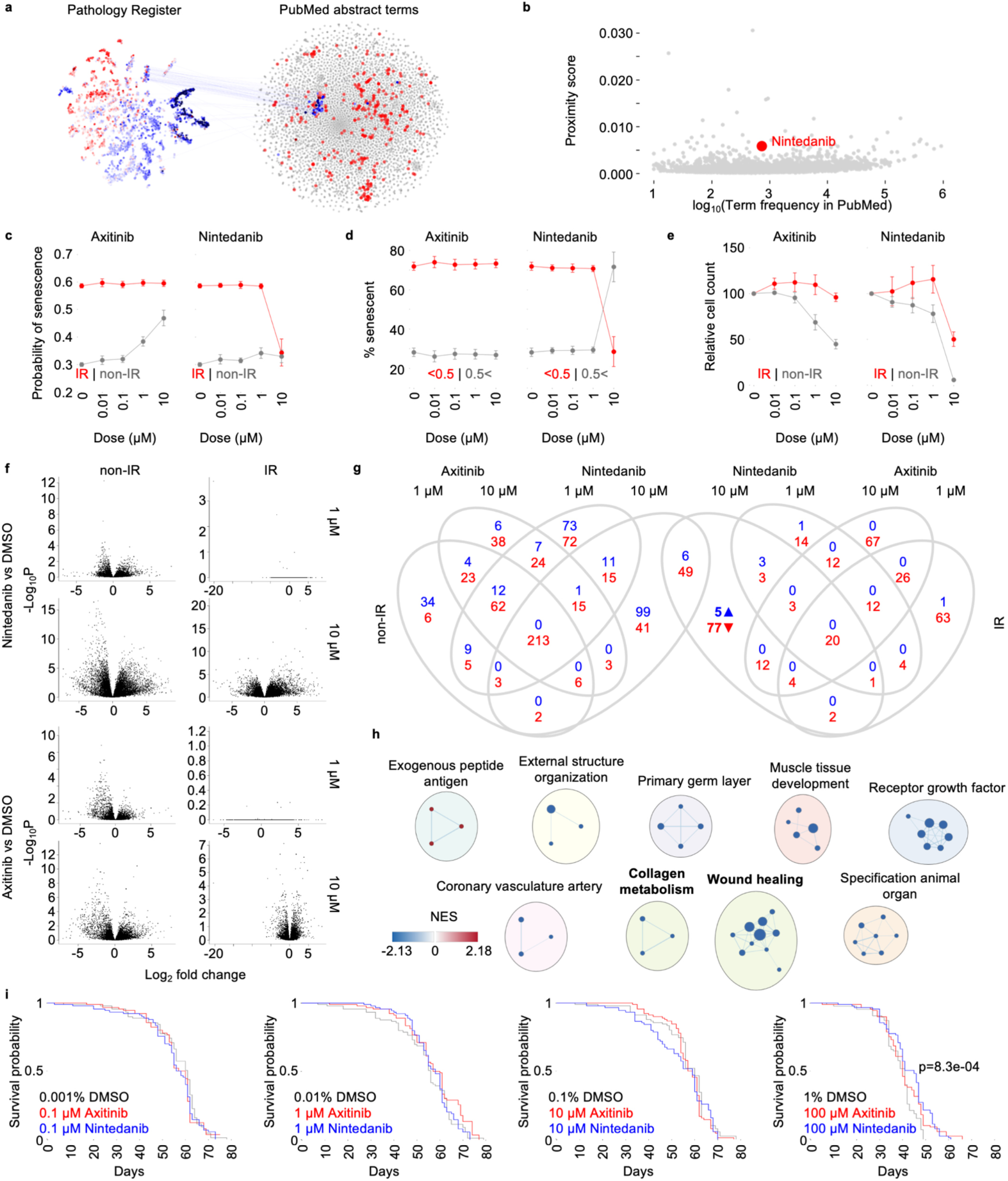
Nintedanib reduces senescence in cells in culture and extends the lifespan of fruit flies. **a**, Lung aging terms from pathology register combined with molecules in PubMed abstracts. **b**, Proximity scoring of candidate compounds. **c**, Relative cell count (normalized to no drug treatment control) in both ionizing radiation (IR) and non-IR plates (n=3, mean mean ± SEM). **d**, Percentage of IR exposed cells predicted to be senescent above 0.5 threshold (n=3, mean mean ± SEM). **e**, Predicted (deep neural network) probability of senescence IR and non-IR plates (n=3, mean mean ± SEM). **f**, RNA seq volcano plots of respective enrichment analyses (n=3). **g**, Venn diagram showing common significantly enriched pathways (GSEA) at FDR<0.05 confidence between each case and the respective control DMSO. **h**, EnrichmentMap showing clusters of significantly enriched pathways (GSEA). **i**, Drosophila melanogaster survival curves (n=90).

### Nintedanib reduces cellular senescence and extends the lifespan of fruit flies

To explore whether nintedanib could impact aging we tested the effect of the drug on cellular senescence, a cellular model of aging that has been implicated in lung fibrosis^14^. We induced senescence in human dermal fibroblasts by ionizing radiation (IR) exposure and used our recently published senescence predictor^15^ to explore the effect on the cells. Interestingly, 10 µM dose of nintedanib reduced predicted senescence in IR induced senescent fibroblasts (Fig. 4c,d). However, we also observed a cytotoxic effect with a 10 µM dose of nintedanib in both IR and non-IR exposed cells manifested in a significant decrease in the relative cell count (Fig. 4e).

To further understand the effect of nintedanib on senescent cells, we explored changes in global gene expression through RNA-seq (Fig. 4f). Since nintedanib and axinitinb share common targets^16^, we isolated pathways (Fig. 4g) which were changed only in senescent cells treated with nintedanib. Notably, nintedanib downregulated (Fig. 4h) collagen metabolic processes and wound healing gene pathways which have both been implicated in lung fibrosis and aging^17,18^.

To explore whether nintedanib could impact aging *in vivo* we investigated the effect of the drug on the life- and health span of the common aging model organism *Drosophila melanogaster*, specifically the wild-type *w*^*1118*^ fly. We observed a significant increase in the maximum lifespan of fruit flies fed a diet containing 100 µM dose of nintedanib compared to dimethyl sulfoxide (DMSO) vehicle (Fig. 4i). Notably, the increase in lifespan is observed in late life. In total, the human pathome allowed us to identify a drug that may affect the aging process.

## Discussion

In this paper we present the human aging pathome (pathoage.com), a compendium of tissue-specific age- and mortality-associated clinical features extracted from the clinical text narratives in The Danish Pathology Register. Our investigation shows strong age-related variance along two trajectories fitting with the ages of development^19^ and with pathological aging^3^. Strikingly, we observed sex-specific differences in the onset and rate of aging related changes. In males, we observed an onset of aging related changes around forty years of age, while in females, we observe that aging trajectories occur almost immediately after development. This could be considered as evidence towards the hypothesis that aging can be a selected trait in evolution since it occurs in women prior to peak fertility. Although speculative, these findings could also suggest that evolution may have allowed successful males to age later perhaps allowing greater reproduction. It is notable, that patterns of aging in males occur around the time of mean life-expectancy of ancient man^20^.

We assessed whether age could be predicted from clinical text features in lung records and found relatively poor predictive power considering individual features. This is not entirely surprising given the abstract nature of language. Nonetheless, even relatively poor predictive power can reveal useful patterns with the terms ‘carcinoma’^21^ and ‘sarcoidosis’^22^ ranked among the most important to prediction accuracy. To improve accuracy, we investigated whether a collection of associated terms (topics) could contribute to greater predictive power. Indeed, the predicative power of the topics was approximately 2-fold greater than that of any individual term and pathology records closely associated with the lung aging topic showed strong predicative power.

As an example of the utility of the Pathome, we mined PubMed abstracts for molecules that occur frequently together with aging lung terms from the pathology register and identified nintedanib as a potential drug affecting aging. Our investigation of global gene changes shows that nintedanib down-regulates collagen metabolism and wound healing pathways in senescent human dermal fibroblasts.

This is compatible with evidence that idiopathic pulmonary fibrosis is characterized by the accumulation of collagen^18^ and an altered wound healing in response to persistent lung injury^23^. It is important to highlight that the method used to identify nintedanib can be used to identify any term associated with aging such as the discovery of new genetic components of aging. The method can also be applied to identify concepts associated with any pathology described in the database. For instance, drugs that may impact liver fibrosis, neurodegeneration or any other defined pathology can be explored.

In sum, our investigation revealed population-level patterns of aging that are connected with developmental and pathological aging. This allows us to identify modifiers of aging that can be translated into new aging interventions. Lastly, we present The Human Pathome, a unique compendium of thousands of tissue-specific aging and mortality associated features.

## Methods

### Danish dictionary of clinical terms

To help identify clinical features in the pathology register we constructed a dictionary of clinical terms in Danish from the patoSnoMed ontology (www.patobank/snomed) and the Danish version of the Systematized Nomenclature of Medicine — Clinical Terms (SNOMED CT)^24^ ontology (https://sundhedsdatastyrelsen.dk/snomedct). Terms found in these ontologies may consist of several words (ex. ‘severe inflammation’). In addition to using such multi-word terms, we added individual words from multi word terms to our dictionary (ex. ‘severe’ and ‘inflammation’.)

### Clinical term extraction

We identified a total of 2,665,283 unique terms, 178,226 unigrams (one word) and 2,487,957 bigrams (two consecutive words) in 32,961,459 pathology text records in The Danish National Pathology Register. This yielded a binary matrix of 32,961,459 samples and 2,665,283 features. We filtered this initial dataset keeping only terms that exist in our dictionary of Danish clinical terms, reducing the size of the dataset to 20,316,270 records and 16,237 terms. We kept terms that appeared at least 50 times in the entire pathology register, and records with 5 or more features present. We then created individual datasets for tissue specific records. We identified records associated with specific tissues using a topology (T) code assigned to each record in the register. For example, to construct a dataset of skeletal system tissues (T10000) we collected all tissues assigned with a topology code that begins with ‘T1’ thereby also including bone tissue (T11000). We applied the same filtering strategy used in the entire dataset to tissue specific datasets. We extracted 242,284 records and 4,684 terms for skeletal tissue (T10000), 177,795 records and 4,275 terms for lung (T28000) and 156,057 records and 4,048 terms for liver (T56000) among other tissues.

### Term normalization

We normalized the clinical term matrix to a term frequency–inverse document frequency (tf-idf) representation. The tf-idf representation for a term t in a document d in a document set consisting of n documents is tf-idf(t,d)=tf(t,d)*idf(t), tf(t,d) being the frequency of a term t in document d, and and idf being idf(t)=log[n/df(t)]+1 (df(t) is the frequency of term t in all documents in the document set). To identify the average term frequency within each age-group we calculated the mean value of all record vectors within each age group. This yielded one term vector per age group. We calculated the mean incidence age of clinical terms in the entire register and in each of the tissue specific datasets.

### Topic modeling with Latent Dirichlet allocation (LDA)

We used the scikit-learn implementation of Latent Dirichlet allocation (LDA)^9^ to identify latent semantic structures in the entire corpus of records within tissue specific datasets. We ran LDA using the batch variational Bayes method. To determine the optimal number of topics yielding the best fit for the model we employed a perplexity minimization approach^25^ by repeatedly fitting an LDA model to our dataset and varying the number of topics (2-140). We then identified the number of topics associated with the smallest perplexity score to be optimal. The topic model yields topic word distributions signifying the number of times each word is assigned to a topic. Similarly, the topic model also yields document topic distribution signifying the degree to which a topic is associated with a document.

### Age-prediction from clinical features

We used a deep neural network (DNN) multi-layer perceptron (MLP) regression to predict age from clinical text features in the one-hot representation of the clinical term matrix. We then calculated the model coefficient of determination score (R^2^) and the median absolute error (MAE) for each model. We used ordinary least squares (OLS) linear regression to regress the predicted age and chronological age of records and calculated topic specific regression slopes. We used the scikit-learn permutation_importance function to inspect our DNN age-prediction model to assess the impact of individual features on the model’s accuracy measured by the model’s coefficient of determination score (R^2^).

### Permutation topic importance

To assess the impact of a collection of associated terms (topic) on the model’s age-prediction accuracy we shuffled the collected term vectors within a topic and calculated the change in the model coefficient of determination score (R^2^). Topics associated with a greater change are deemed more important to age-prediction.

### Dimensionality reduction

We used the scikit-learn implementation of PCA and t-SNE. We applied PCA and t-SNE to age-aggregated tf-idf term matrices. We used the python umap-learn package implementation of UMAP on tf-idf term matrices.

### Term enrichment in tissue and morphology-specific records

Term enrichment in tissue or morphology specific records is calculated as log((B+1)/(A-B+1)) where B is the frequency of a term in tissue or morphology specific records and A is the frequency a term in the entire dataset.

### PubMed term proximity score

We extracted a total of 175,555 unique terms from 31,850,051 PubMed abstracts. Given a binary feature matrix M and a set A of terms within matrix M we calculated a proximity score for each individual term in a given set B within matrix M. We applied a tf-idf transformation to the feature matrix M and calculated the cosine distances between individual terms in set A to individual terms in set B yielding a distance matrix AxB. We calculated the term proximity score to be the mean distance of each term b in set B to all terms in set A that are co-mentioned with term b at least once.

Matrix M: PubMed abstracts years 2000 onwards.

Set A: Aging lung terms.

Set B: All PubChem compounds that occur in PubMed abstracts ten times or more.

### Term and topic associated mortality

For term and topic associated mortality, we used the R survival package to perform Cox survival regression. For each clinical term we calculated a Cox regression coefficient reflecting the hazard associated with the incidence of the term in pathology records, adjusted for word count and birth year cohort. For topic-associated mortality, we stratified patient pathology records according to topics. We created a Boolean variable for each topic reflecting the association of a pathology record with a topic. This yielded a matrix of records and topics. We performed Cox survival regression on the time to death from examination and noted the Cox regression coefficient associated with each topic reflecting the hazard associated with the incidence of the topic in pathology records.

### Cell culture

Human primary fibroblast cell lines (Coriell, NJ, USA) AG08498 (AG), GM22159 (159) and GM22222 (222) were cultured in 4.5g/L-enriched Dulbecco’s Modified Eagle’s Medium (DMEM)/ Ham’s F-12 Nutrient Mix (F12) in a 1:1 solution supplemented with 10% fetal bovine serum (FBS) and 1% penicillin/streptomycin. Cells were maintained at 37°C in 5% CO2 atmosphere conditions and passaged every 2-3 days. For senescence assays, cells at 70-80% confluency and below 20 passages were seeded in 96-well plates (Corning, 3340) at a density of 3000 cells/well and incubated overnight at 37 °C and 5% CO2. Control plates were seeded at 3000 cells/well or 1500 cells/well. One day after seeding, plates were irradiated using a YXLON Smart Maxi Shot. Cells were exposed with emission of 0.85 Gy/min. for 12 minutes for a total exposure of 10 Gy. After IR exposure, cells were incubated for 6 days with medium changed every 48h. Control plates were seeded on day 6. On day 7 cells were treated with compounds or vehicle for 48 hours after which the cells were either harvested for RNA or fixed with 4% paraformaldehyde for 10 min, washed in PBS and stained with DAPI. Cells were subsequently imaged using an IN Cell analyzer 2200 high content microscopy at 20x magnification, 12 fields per well.

### RNA sequencing

RNA was extracted using Trizol according to manufacturer’s protocol. DNBSEQ Eukaryotic Long Non-Coding RNA-sequencing was performed by BGI Denmark. Mapping-based quantification of the GRCh38 transcriptome from RNA sequencing paired-end reads was performed with salmon^26^ using a pre-computed transcriptome index for salmon obtained from refgenie^27^.Differential expression analysis was performed with DESeq2 1.38.2^28^ on genes mapped from transcripts with the gencode annotation of the Ensembl gene set downloaded from refgenie http://refgenomes.databio.org/v3/assets/splash/2230c535660fb4774114bfa966a62f8 23fdb6d21acf138d4/salmon_sa_index?tag=default. Genes with fewer than 10 reads across all samples were filtered prior to all downstream analyses. Gene set enrichment analysis was performed using GSEA 4.3.2^29^. An expression dataset file (.gct) was prepared using DESeq2^28^ normalized counts for all samples. Phenotype labels files (.cls) were prepared for each of the group comparisons. GSEA was run on with the gene set database ‘MSigDB c5.go.bp.v2022.1.Hs.symbols.gmt’ ^30^and gene_set permutation type. We used gene sets which are significantly enriched (upregulated) at FDR<0.05 in each phenotype in all downstream analyses. Significantly enriched pathways from GSEA^29^ were visualized in Cytoscape^31^ using the EnrichmentMap, AutoAnnotate, WordCloud and clusterMaker2 applications.

### Fruit fly maintenance

All diets were made on a standard diet (SD) base consisting of 47.5 g cornflour, 41.6 g dextrose, 19.3 g Brewer’s Yeast, 6.55 g Low Melting Agar (Calbiochem), and 2.46% Nipagin (Merck, Germany) per litre. All ingredients except Nipagin were mixed and heated to 80°C. When the mixture had cooled to 40°C, Nipagin was added. The mix was distributed in falcon tubes and compounds added in various concentrations to make the treatment diets. Diets with equivalent amounts of DMSO were used as controls. Stock flies were housed in vials of 30 flies to avoid overcrowding and kept on the standard diet. Both stock and treatment flies were kept at a constant temperature of 25°C, a relative humidity of 60%, and a 12:12 h light:dark cycle. The wild type strain *w*^*1118*^ (Bloomington Drosophila Stock Center) was used for all longevity assay. Before assays, 5-10 crosses with a ratio of 15:9 female to male flies were set and kept under standard rearing conditions in polypropylene vials in standard diet.

### Fruit fly lifespan assay

Every three days, for 9-12 days, flies were flipped into new vials containing the standard diet. Hatches were collected at birth and put in new vials with the desired compound condition. Per each condition, three vials with ten male flies each were prepared. Vials were put in front of cameras for our fly tracking system as part of the Tracked.bio platform (www.tracked.bio). For all longevity assay, flies were flipped once weekly into vials with freshly prepared food. Each vial had ten male flies, which were selected from the new-born hatches of the set crosses. Male flies were chosen among those which did not show any damage to the wings.

During each flipping, flies were counted, and data collected into a spreadsheet. Behavioral metrics were calculated from the Tracked.bio system. We used the lifelines python package to fit a Kaplan-Meier estimator for the survival function of fruit fly lifespan and to perform a log-rank test to test for statistically significant differences in survival. A count of live flies in vials was recorded once per week. Since fruit fly vials were initiated over a period of several days as newly hatched flies were collected, we extrapolated weekly counts to daily counts before performing survival analysis.

## Acknowledgements

This research was supported by the Novo Nordisk Foundation Challenge Programme (#NNF17OC0027812), the Nordea Foundation (#02-2017-1749), the Neye Foundation, the Lundbeck Foundation (#R324-2019-1492), the Ministry of Higher Education and Science (#0238-00003B) and Insilico Medicine. The funders had no role in study design, data collection and analysis, decision to publish or preparation of the manuscript.

## Figure Legends

**Extended Data Fig. 1** | **Sex-specific patterns. a**, UMAP of clinical features in pathology records from males in the entire pathology register. **b**, UMAP of clinical features in pathology records from females in the entire pathology register. **c**, t-SNE of clinical features in age-aggregated pathology records from males in the entire pathology register. **d**, t-SNE of clinical features in age-aggregated pathology records from females in the entire pathology register. **e**, Positive enrichment of clinical terms in various tissue and morphology specific records **f**, Term count associated hazard. **g**, Birth cohort associated hazard.

**Extended Data Fig. 2** | **Tissue-specific analyses. a**, PCA of age-aggregated tissues specific pathology records. **b**, UMAP of clinical features of tissue specific pathology records (mean incidence age). **c**, UMAP of clinical features of tissue specific pathology records (Cox regression coefficient). **d**, Normalized Euclidean distance between age adjacent PCA coordinates of all tissues in.

**Extended Data Fig. 3** | **Topic modelling. a**, LDA perplexity for model fitted with varying number of topics (skeletal system, lung, liver). **b**, Topic associated mean age. **c**, Topic associated mortality. **d**, Topic associated age-prediction linear regression. **e**, Topic associated permutation feature importance (PFI). **f**, Topic associated age-prediction regression slope. **g**, Terms describing Human Immunodeficiency Virus (HIV). **h**, Bivariate kernel density estimation (kde) and histograms of chronological age vs. predicted age, records closely associated with HIV and t-SNE of clinical feature LDA distributions in topics highlighting OGPC, PA and HIV topics.

