## Extended data for "The human pathome shows sex specific aging patterns post-development"

Contains:

- Figure legends for Extended Data Figures 1-3.
- Extended Data Figures 1-3.

### Extended Data Figure Legends

**Extended Data Fig. 1 | Sex-specific patterns. a,** UMAP of clinical features in pathology records from males in the entire pathology register. **b,** UMAP of clinical features in pathology records from females in the entire pathology register. **c,** t-SNE of clinical features in age-aggregated pathology records from males in the entire pathology register. **d,** t-SNE of clinical features in age-aggregated pathology records from females in the entire pathology register. **e,** Positive enrichment of clinical terms in various tissue and morphology specific records **f,** Term count associated hazard. **g,** Birth cohort associated hazard.

**Extended Data Fig. 2 | Tissue-specific analyses. a,** PCA of age-aggregated tissues specific pathology records. **b,** UMAP of clinical features of tissue specific pathology records (mean incidence age). **c,** UMAP of clinical features of tissue specific pathology records (Cox regression coefficient). **d,** Normalized Euclidean distance between age adjacent PCA coordinates of all tissues in.

**Extended Data Fig. 3 | Topic modelling. a,** LDA perplexity for model fitted with varying number of topics (skeletal system, lung, liver). **b,** Topic associated mean age. **c,** Topic associated mortality. **d,** Topic associated age-prediction linear regression. **e,** Topic associated permutation feature importance (PFI). **f,** Topic associated age-prediction regression slope. **g,** Terms describing Human Immunodeficiency Virus (HIV). **h,** Bivariate kernel density estimation (kde) and histograms of chronological age vs.

predicted age, records closely associated with HIV and t-SNE of clinical feature LDA distributions in topics highlighting OGPC, PA and HIV topics.

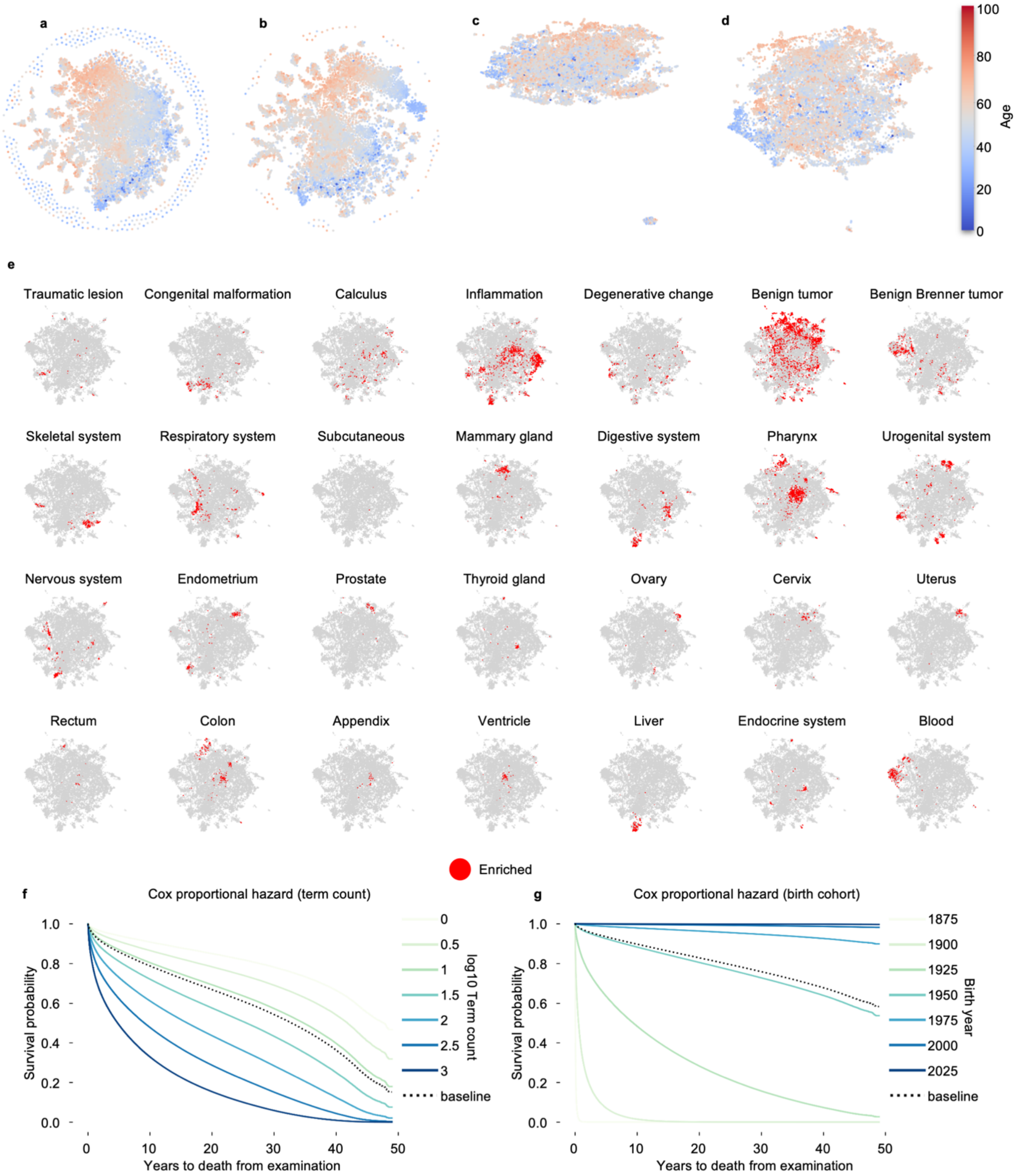


**Extended Data Fig. 1 | Sex-specific patterns. a,** UMAP of clinical features in pathology records from males in the entire pathology register. **b,** UMAP of clinical features in pathology records from females in the entire pathology register. **c,** t-SNE of clinical features in age-aggregated pathology records from males in the entire pathology register. **d,** t-SNE of clinical features in age-aggregated pathology records from females in the entire pathology register. **e,** Positive enrichment of clinical terms in various tissue and morphology specific records **f,** Term count associated hazard. **g,** Birth cohort associated hazard.

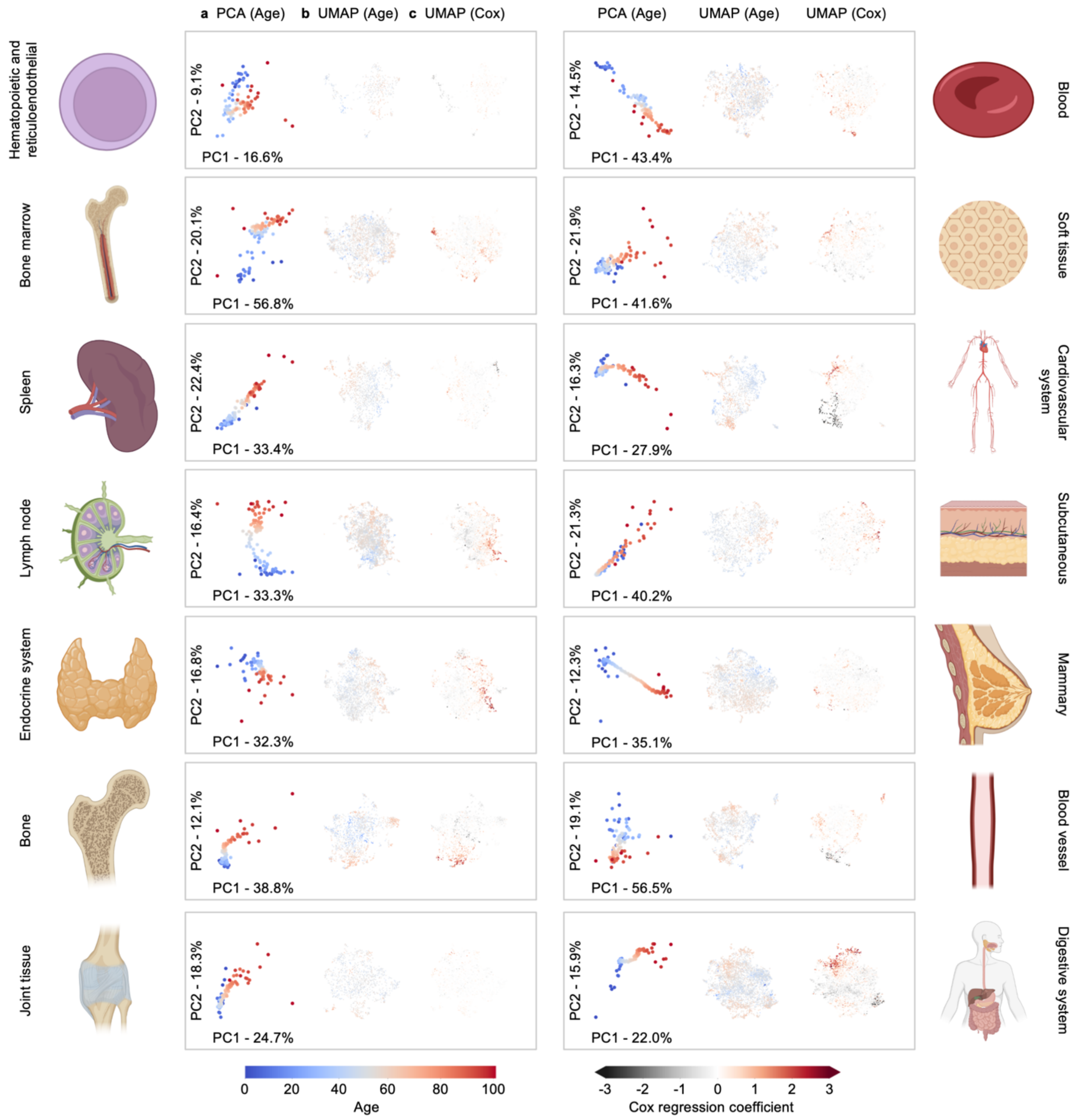


**Extended Data Fig. 2 | Tissue-specific analyses. a,** PCA of age-aggregated tissues specific pathology records. **b,** UMAP of clinical features of tissue specific pathology records (mean incidence age). **c,** UMAP of clinical features of tissue specific pathology records (Cox regression coefficient). **d,** Normalized Euclidean distance between age adjacent PCA coordinates of all tissues in.

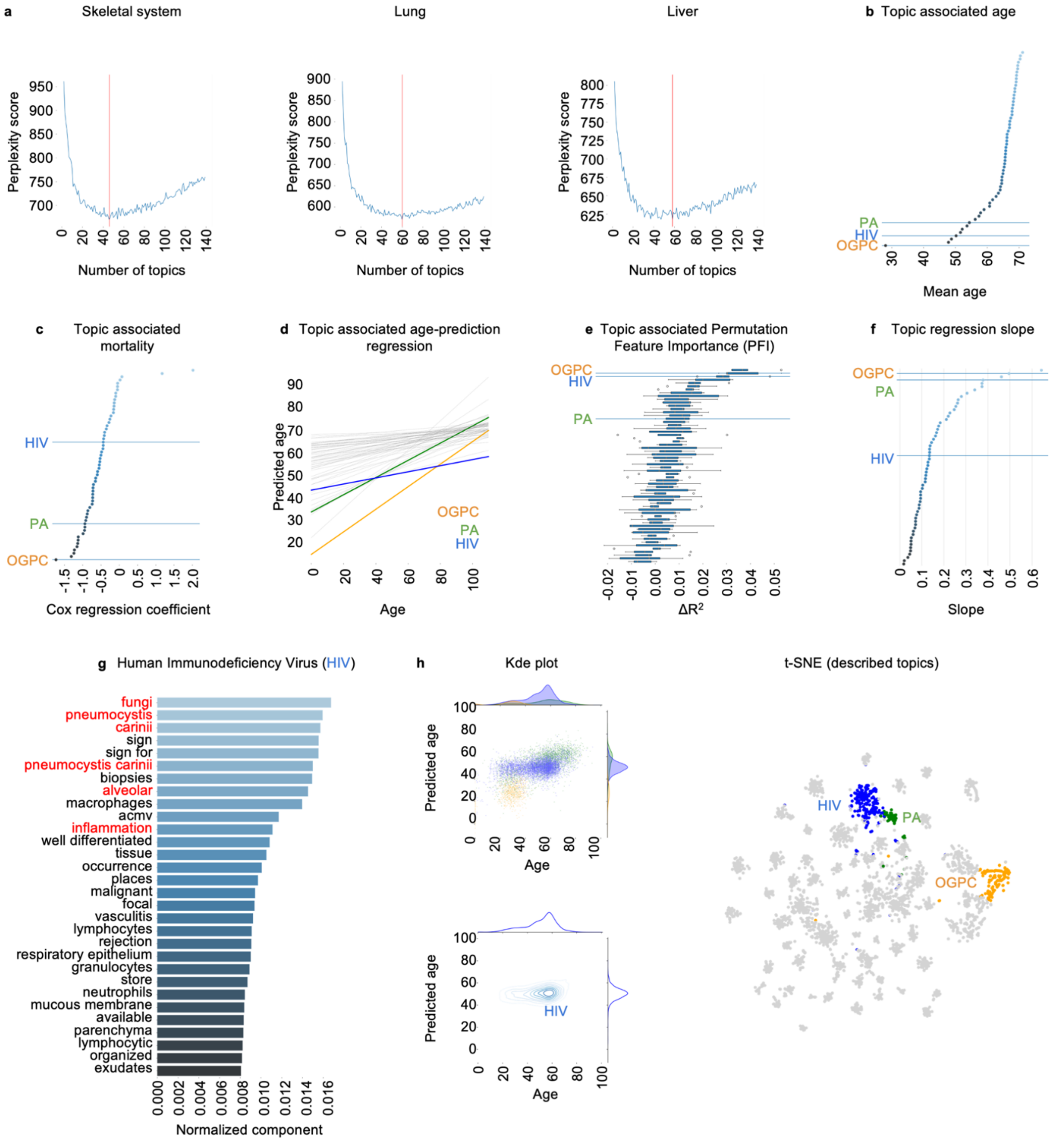


**Extended Data Fig. 3 | Topic modelling. a,** LDA perplexity for model fitted with varying number of topics (skeletal system, lung, liver). **b,** Topic associated mean age. **c,** Topic associated mortality. **d,** Topic associated age-prediction linear regression. **e,** Topic associated permutation feature importance (PFI). **f,** Topic associated age-prediction regression slope. **g,** Terms describing Human Immunodeficiency Virus (HIV). **h,** Bivariate kernel density estimation (kde) and histograms of chronological age vs. predicted age, records closely associated with HIV and t-SNE of clinical feature LDA distributions in topics highlighting OGPC, PA and HIV topics.
